# Selective Expression of Variant Surface Antigens Enables *Plasmodium falciparum* to Evade Immune Clearance *in vivo*

**DOI:** 10.1101/2020.07.22.215640

**Authors:** Marvin Chew, Weijian Ye, Radoslaw Igor Omelianczyk, Charisse Flerida Pasaje, Regina Hoo, Qingfeng Chen, Jacquin C. Niles, Jianzhu Chen, Peter Preiser

## Abstract

*Plasmodium falciparum* has developed extensive mechanisms to evade host immune clearance. Currently, most of our understanding is based on *in vitro* studies of individual parasite variant surface antigens and how this relates to the processes *in vivo* is not well-understood. Here, we have used a humanized mouse model to identify parasite factors important for *in vivo* growth. We show that upregulation of the specific PfEMP1, VAR2CSA and the RIFIN PF3D7_1254800 provides the parasite with protection from macrophage phagocytosis and natural killer cell mediated killing. Taken together, these findings reveal new insights on the molecular and cellular mechanisms that coordinate the immune escape process the parasite utilizes *in vivo*. As immune evasion may be particularly important during the establishment of the blood stage infection when parasite numbers are still relatively small, identification of specific parasite variant surface antigens provides targets for developing more effective vaccines by targeting parasite immune evasion.

## Introduction

*Plasmodium falciparum* is the most important causative agent of human malaria. Currently, annual malaria infections and mortality are approximately 220 million and 400,000, respectively (Gross, 2019). To control malaria infection in such a large scale, an effective vaccine is essential. However, although a pre-erythrocytic stage vaccine for malaria, Mosquirix, has been approved, its efficacy is only ∼36% (Theander and Lusingu, 2015). To further improve the efficacy of malaria vaccine, the incorporation of blood-stage antigens into a multistage vaccine has been suggested (Miura, 2016). Many studies have suggested that antibodies against surface-exposed antigens, such as the PfEMP1s encoded by *var* genes and RIFINs encoded by *rif* genes can confer some degree of clinical protection (Abdel-Latif et al., 2002; Chan et al., 2012). However, given the large diversity in these variant surface antigens (VSA), no protective epitope has been identified as yet (Dodoo et al., 2001; Mackintosh et al., 2008; Makobongo et al., 2006).

Macrophages and natural killer (NK) cells are the earliest innate immune cells that respond to parasite infection (Chen et al., 2014; Lai et al., 2018), and the outcome of this early host– parasite interaction is a strong determinant for immunopathology and disease severity (Urban et al., 2005; Ye et al., 2018). However, parasites have developed mechanisms to inhibit macrophage phagocytosis (Gomes et al., 2016; Serghides et al., 2006) and evade killing by NK cells (Saito et al., 2017; Ye et al., 2018). Macrophages recognize and phagocytose parasite-infected red blood cells (iRBC) through the class B scavenger receptor, CD36, which binds to group B and C PfEMP1 expressed on the surface of iRBC (Patel et al., 2004). Parasites isolated from patients with severe malaria exhibit reduced expression of PfEMP1 that binds to CD36 (Bernabeu et al., 2016). A subset of RIFIN have also been shown to inhibit NK cell activation through leukocyte immunoglobulin-like receptor subfamily B member 1 (LILRB1) (Saito et al., 2017). Despite the progress, the determinants that allow a parasite to evade macrophage and NK cell clearance and thrive in the human host have not been identified. It is therefore critical for the development of more effective vaccines to take a broader immunological approach that also considers antigens that facilitate immune escape (Rénia and Goh, 2016).

One of the main challenges of immunological studies of *P. falciparum* is the lack of a suitable model system that accurately emulates all the intricacies of the complex immune responses the human host develops in response to an infection. Current studies using *in vitro* cultured parasites are further hampered by the fact that through continuous *in vitro* culture, numerous parasitic adaptations have occurred, including the loss of parasitic virulence factors (Bull et al., 1991; Day et al., 1993; Langreth and Peterson, 1985; Udeinya et al., 1983). Noticeably, there is an overall downregulation of PfEMP1 (Peters et al., 2007) and specific downregulation of the non-CD36 binding Group A *var* genes (Janes et al., 2011). Interestingly, parasites isolated from patients with severe malaria expressed higher level of group A *var* transcripts compared to parasites isolated from patients with non-severe malaria (Bernabeu et al., 2016). Similarly, expression of RIFIN as well as STEVOR, two additional families of variant surface antigens, are absent or expressed at a lower level in continous *in vitro* cultured strains compared to parasites directly isolated from patients (Fernandez et al., 1999) (Blythe et al., 2008).

To gain a better understanding in the mechanisms the parasite utilizes to effectively survive *in vivo* we characterized the changes that occur when *in vitro* cultured *P. falciparum* is adapted to grow in human RBC-reconstituted NOD/SCID IL2γ^null^ (RBC-NSG) mice. In RBC-NSG mice, human RBCs support parasite infection in the presence of mouse macrophages as NSG mice are deficient in T, B and natural killer (NK) cells. The system provides a reductionist *in vivo* environment to study the changes the parasites need to undergo to evade this first-line of defense of the innate immune system. We show that in the presence of physical and immune stress, parasites upregulate the expression of specific *var* and *rif*. Specifically, we identify that upregulation of VAR2CSA, a member of PfEMP1 that does not bind to CD36, enables parasites to escape from macrophage phagocytosis while upregulation of RIFIN encoded by PF3D7_1254800 inhibits NK cell-mediated clearance through inhibitory LILRB1. Our findings reveal molecular and cellular mechanisms involved in parasite adaptation to immunological stress. As escape from clearance by the innate immune system is particularly important early on during the establishment of the blood stage infection when parasite numbers are still relatively small, targeting the select PfEMP1 and RIFIN may lead to improved intervention strategies against malaria infection.

## Results

### Patent infection of RBC-NSG mice by *P. falciparum* requires a period of adaptation

To study *P. falciparum* adaptation to immunological and physical stresses *in vivo*, we infected RBC-NSG mice with an equal inoculum of six different *P. falciparum* strains that had been continuously cultured *in vitro* (Table 1). In all cases, no parasites were detected by Giemsa staining of peripheral blood smears until at least day 18 after the initial inoculation (Figure 1A and data not shown), consistent with a previous report (Angulo-Barturen et al., 2008). By 35 days after the initial inoculation when parasitemia was apparent, blood was collected from the infected mice and used to infect human RBCs *in vitro*. After at least 20 cycles of expansion, the recovered parasites were used to infect a new batch of RBC-NSG mice. No delay in parasitemia was observed (Figure 1B). These results show that parasites undergo adaptation in RBC-NSG mice to acquire a phenotype that is stably maintained.

**Table 1.** List of parasites adapted and used

| Parasite | Source | Description | Day post infection for adapted parasite to appear in peripheral blood |
| --- | --- | --- | --- |
| 3D7 | BEI Resources (MRA-102) | Clone derived from NF54 by limiting dilution (Walliker et al., 1987). | 3D7 B7 AS – Day 27 |
| W2mef | BEI Resources (MRA-615) | Mefloquine-resistant culture lines derived from Indochina strain (Oduola et al., 1988). | W2mef AS – Day 19 |
| T994 | BEI Resources (MRA-153) | Isolate derived from T9 strain collected from patient in Thailand (Fenton et al., 1989). | T994 AS – Day 21 |
| NF54attB | Niles lab | NF54attB that harbors an attB site at the Cg6 locus on Chr 7 (Adjalley et al., 2011). | NF54attB AS – Day 24 |
| 3D7attB | BEI Resources (MRA-845) | 3D7 parasite that harbors an attB site at the Cg6 locus on Chr 7 (Nkrumah et al., 2006). | 3D7attB AS – Day 30 |
| NF54attB<sup>Cas9+T7</sup><br>Polymerase (NF54CR) | Niles lab | NF54 parasites that expressed Cas9 and T7 RNA polymerase (Sidik et al., 2016). | NF54CR AS – Day 25 |
| NF54CR AS<br>*mahrp1* | This paper | *mahrp1* gene under the regulation of aTc on the NF54CR AS background. | Adapted parasites used for transfection |
| 3D7-SLI-RIFIN* | This paper | 3D7 strain with PF3D7_1254800 gene tagged by a resistant marker for continuous expression. | N.A. |

\* Not adapted

**Figure 1.**
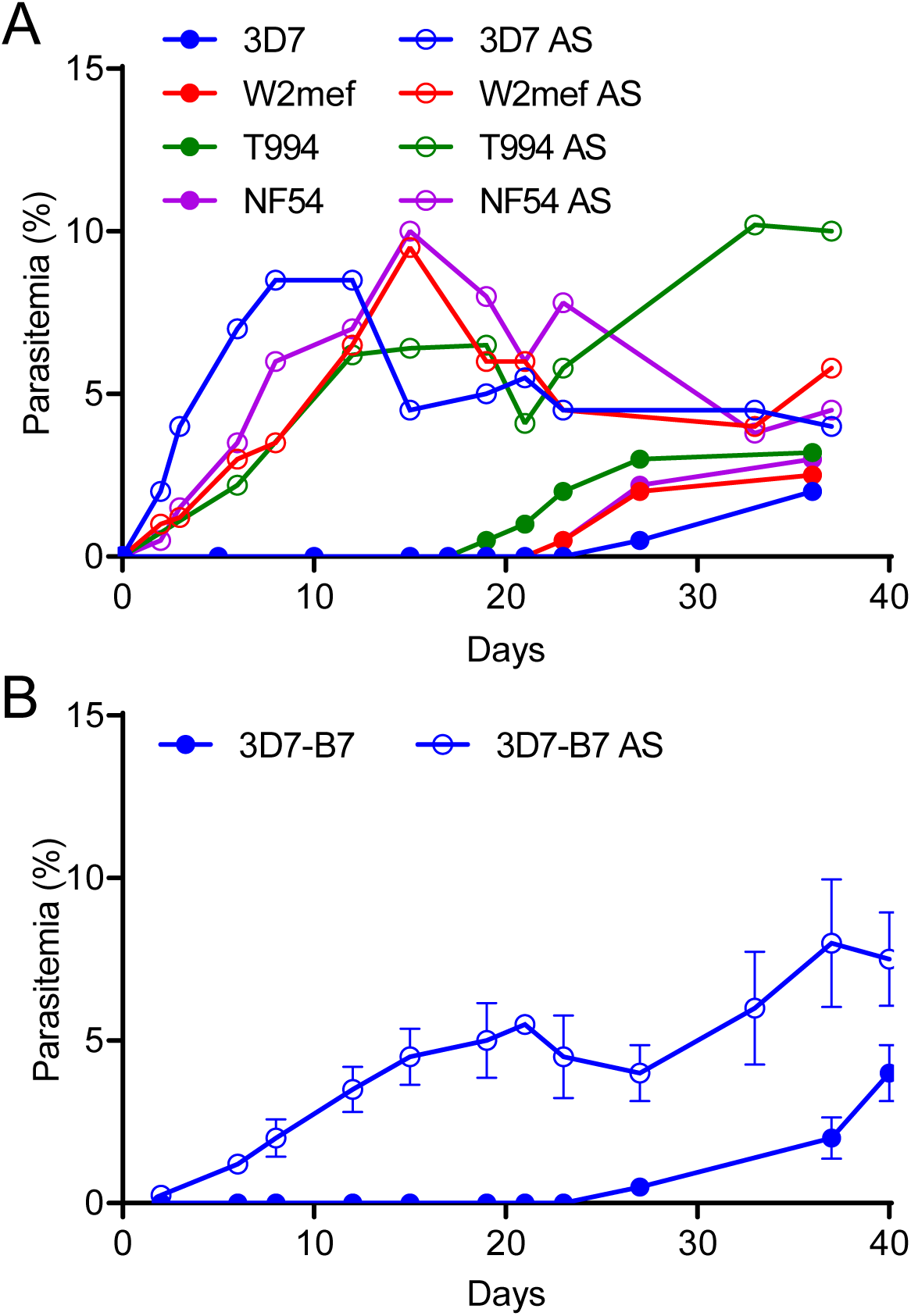
Comparison of parasitemia of non-adapted and adapted *P. falciparum* strains in RBC-NSG mice. **(A** and **B)** RBC-NSG mice were infected with the indicated strains of non-adapted (**A**) or adapted (AS) (**B**)*P. falciparum* strains, and parasitemia was assessed by Giemsa staining of peripheral blood smears. Each point represents the mean of three independent biological repeats. **(C)** A single clone of 3D7, 3D7-B7, was isolated, expanded and used to infect RBC-NSG mice. After 40 days, parasites from peripheral blood was recovered, cultured and expanded in vitro to generate 3D7-B7 AS. 3D7-B7 AS parasites were then used to infect a new batch of RBC-NSG mice. Each point represents the mean ± SEM. of three independent biological repeats.

Since long-term continuous *in vitro* culture of *P. falciparum* leads to genetic heterogeneity (Yeda et al., 2016), we serially diluted the non-adapted 3D7 strain of *P. falciparum* to obtain a single clone, 3D7-B7, and repeated the adaptation experiment in RBC-NSG mice. Again there was a significant delay of 26 ± 5.1 days before 3D7-B7 was detected in the blood of inoculated mice (Figure 1C). When the recovered parasites from RBC-NSG mice were cultured *in vitro* for 20 cycles and used to infect RBC-NSG mice, no delay in parasitemia was observed. Thus, the adaptation in RBC-NSG mice is not due to the selection of a single unique parasite from a genetically heterogeneous culture. Parasites recovered from RBC-NSG mice after adaptation are referred to as adapted strains (AS).

### Adaptation of *P. falciparum* parasites is associated with transcriptional changes of specific variant surface antigens

To understand the changes contributing to the adaptation phenotype, we determined the genetic differences between three non-adapted and adapted parasites (3D7-B7, 3D7attB and NF54attB) by whole genome sequencing (Nkrumah et al., 2006). Single nucleotide polymorphism (SNP) variant calling identified similar SNPs in both the parental non-adapted strains and their corresponding adapted strains and no conserved unique SNP was detected among the three adapted strains. Kinship coefficient analysis (Manichaikul et al., 2010) of the SNPs called for each parasite strain pair did not significantly diverge and had high kinship coefficient values (>0.487, Figure 2A). Since kinship coefficient values of above 0.354 corresponds to duplicate or monozygotic twins (Yang et al., 2011), our results suggest that adaptation is not due to the acquisition of a unique mutation(s).

**Figure 2.**
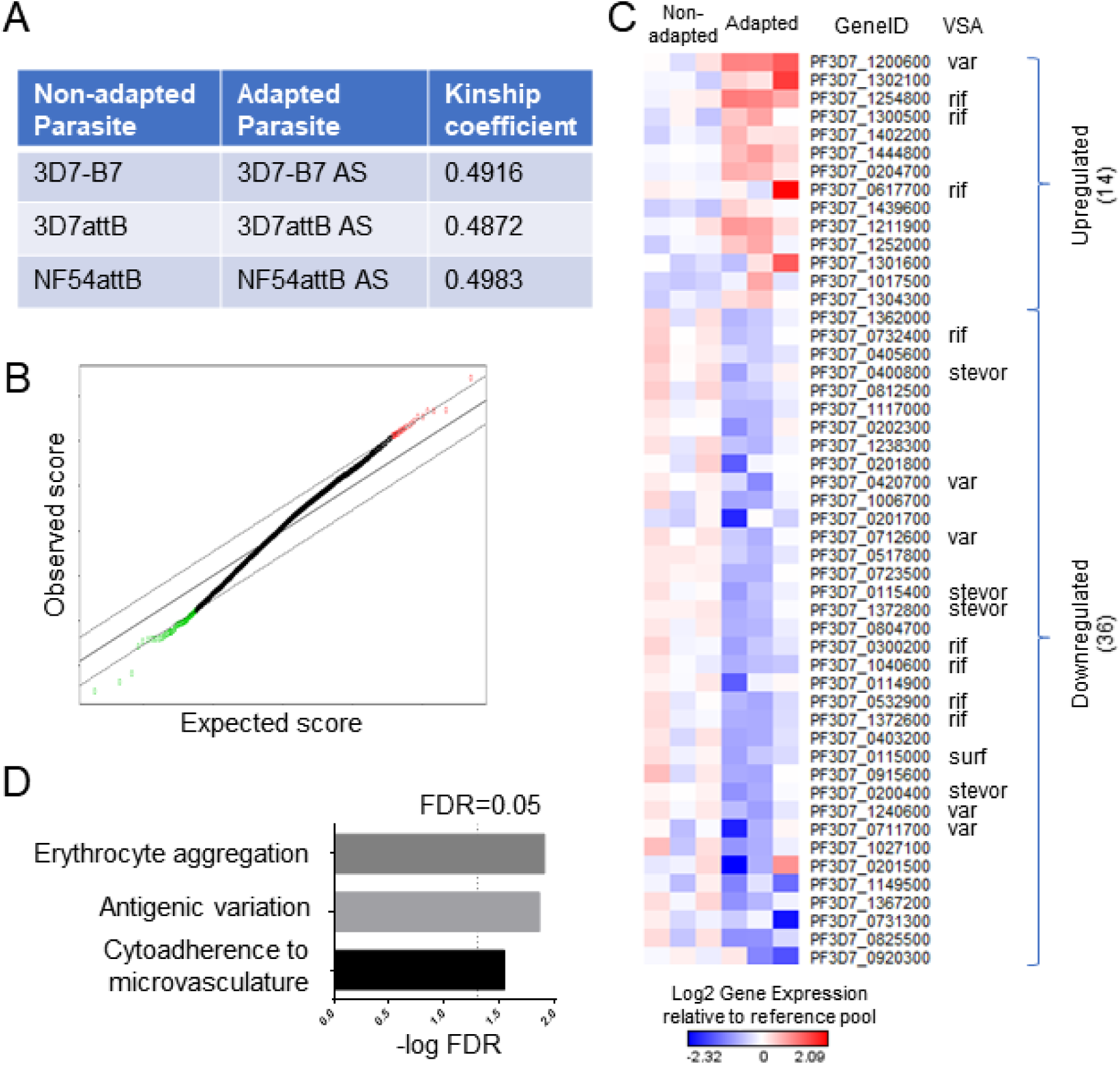
Differences in gene expression between adapted and non-adapted parasites. **(A)** Kinship coefficient analysis of SNPs in adapted and non-adapted parasites. **(B)** SAM of the 3 pairs of non-adapted and adapted parasite strains using a delta of 0.55. Red: upregulated genes; green: downregulated genes. (**C)** Heatmap of upregulated and downregulated DEGs identified using an FDR of 0.01. Each column represents a different parasite strain. Each row represents a DEG. VSAs are identified. **(D)** Top three Go biological processes that were significantly enriched in DEGs.

We next examined whether transcriptional changes could be responsible for the adaptation phenotype by comparing the gene expression profile of three pairs of non-adapted and adapted *P. falciparum* strains (3D7-B7, 3D7attB and NF54CR) using microarray analysis (Bozdech et al., 2003a). We included NF54CR, a strain of NF54attB parasites stably expressing Cas9 and T7 RNA polymerase (Nasamu et al., 2019), as it was used for later studies. Cultures of infected RBCs were tightly synchronized and RNA was harvested every 8 hours across the intraerythrocytic development cycle (IDC) to generate 6 time points per parasite strain. Significance Analysis of Microarray (SAM) (Tusher et al., 2001) of the 3 pairs of non-adapted and adapted parasite strains using a delta of 0.55 showed that 27 genes and 93 genes were significantly upregulated and downregulated, respectively (Figure 2B and S1). Using a false discovery rate (FDR) of 0.01, we further refined these genes to derive a list of differentially expressed genes (DEG), of which 14 were upregulated and 36 were downregulated (Figure 2C and Tables 2 and 3). Gene Ontology (GO) (Ashburner et al., 2000; Consortium, 2018) biological processes analysis (Mi et al., 2018) revealed that the DEGs are involved in processes such as modulation of erythrocyte aggregation, antigenic variation, and cytoadherence to microvasculature (Figure 2D). Many of the DEGs were parasite variant surface antigens, including one *var* gene and 3 *rif* genes that were up-regulated and 4 *var* genes, 5 *rif* genes, 4 *stevor* genes and 1 *surf* gene that were down-regulated (Figure 2C and Tables 2 and 3).

**Table 2.** Upregulated DEGs.

| Gene ID | Description | Bonferroni<br>pval | log2FC |
| --- | --- | --- | --- |
| PF3D7_1200600 | erythrocyte membrane protein 1, PfEMP1 | 5E-16 | 1.139171 |
| PF3D7_1302100 | gamete antigen 27/25 | 3.4E-11 | 0.93061 |
| PF3D7_1254800 | rifin | 2.92E-10 | 0.884545 |
| PF3D7_1300500 | rifin | 2.77E-06 | 0.657873 |
| PF3D7_1402200 | cytochrome c oxidase subunit ApiCOX19,<br>putative | 2.05E-05 | 0.597823 |
| PF3D7_1444800 | fructose-bisphosphate aldolase | 2.37E-05 | 0.593211 |
| PF3D7_0204700 | hexose transporter | 2.72E-05 | 0.5889 |
| PF3D7_0617700 | rifin | 3.14E-05 | 0.584273 |
| PF3D7_1439600 | cytochrome c oxidase subunit ApiCOX26,<br>putative | 3.29E-05 | 0.582841 |
| PF3D7_1211900 | non-SERCA-type Ca<sup>2+</sup> -transporting P-ATPase | 4.41E-05 | 0.573354 |
| PF3D7_1252000 | tRNA Glutamine | 4.77E-05 | 0.57074 |
| PF3D7_1301600 | erythrocyte binding antigen-140 | 4.86E-05 | 0.57017 |
| PF3D7_1017500 | myosin essential light chain ELC | 7.02E-05 | 0.557975 |
| PF3D7_1304300 | conserved Plasmodium protein, unknown<br>function | 9.28E-05 | 0.548597 |

**Table 3.** Downregulated DEGs.

| Gene ID | Description | Bonferroni<br>pval | log2FC |
| --- | --- | --- | --- |
| PF3D7_0920300 | conserved Plasmodium protein, unknown function | 1.29E-08 | -0.79847 |
| PF3D7_0825500 | protein KRI1, putative | 1.48E-08 | -0.79512 |
| PF3D7_0731300 | Plasmodium exported protein (PHISTb), unknown function | 2.12E-08 | -0.7864 |
| PF3D7_1367200 | CLAMP domain-containing protein, putative | 3.45E-08 | -0.77445 |
| PF3D7_1149500 | ring-infected erythrocyte surface antigen 2, pseudogene | 1.78E-07 | -0.7329 |
| PF3D7_0201500 | Plasmodium exported protein (hyp9), unknown function | 4.02E-07 | -0.71138 |
| PF3D7_1027100 | U3 small nucleolar ribonucleoprotein protein MPP10, putative | 5.42E-07 | -0.70335 |
| PF3D7_0711700 | erythrocyte membrane protein 1, PfEMP1 | 6.03E-07 | -0.70048 |
| PF3D7_1240600 | erythrocyte membrane protein 1, PfEMP1 | 9.96E-07 | -0.68674 |
| PF3D7_0200400 | stevor | 1.75E-06 | -0.67099 |
| PF3D7_0915600 | conserved Plasmodium protein, unknown function | 3.62E-06 | -0.65018 |
| PF3D7_0115000 | surface-associated interspersed protein 1.3 (SURFIN 1.3) | 4.33E-06 | -0.64498 |
| PF3D7_0403200 | pre-mRNA splicing factor, putative | 4.83E-06 | -0.64177 |
| PF3D7_1372600 | rifin | 5.69E-06 | -0.6369 |
| PF3D7_0532900 | rifin | 7.11E-06 | -0.63029 |
| PF3D7_0114900 | Plasmodium exported protein, unknown function, pseudogene | 1.48E-05 | -0.60804 |
| PF3D7_1040600 | rifin | 1.51E-05 | -0.60739 |
| PF3D7_0300200 | rifin | 1.64E-05 | -0.60488 |
| PF3D7_0804700 | conserved Plasmodium protein, unknown function | 1.68E-05 | -0.6041 |
| PF3D7_1372800 | stevor | 2.06E-05 | -0.59764 |
| PF3D7_0115400 | stevor | 2.16E-05 | -0.59622 |
| PF3D7_0723500 | dynactin subunit 5, putative | 2.38E-05 | -0.59313 |
| PF3D7_0517800 | apicortin, putative | 3.42E-05 | -0.58156 |
| PF3D7_0712600 | erythrocyte membrane protein 1, PfEMP1 | 3.55E-05 | -0.58037 |
| PF3D7_0201700 | DnaJ protein, putative | 4.09E-05 | -0.57578 |
| PF3D7_1006700 | conserved Plasmodium protein, unknown<br>function | 4.29E-05 | -0.57426 |
| PF3D7_0420700 | erythrocyte membrane protein 1, PfEMP1 | 4.36E-05 | -0.5737 |
| PF3D7_0201800 | knob associated heat shock protein 40 | 4.44E-05 | -0.57314 |
| PF3D7_1238300 | pre-mRNA-splicing factor CWC22, putative | 5.95E-05 | -0.56352 |
| PF3D7_0202300 | Plasmodium exported protein (hyp11), unknown<br>function | 6.44E-05 | -0.56085 |
| PF3D7_1117000 | conserved Plasmodium membrane protein,<br>unknown function | 6.61E-05 | -0.55999 |
| PF3D7_0812500 | RNA-binding protein, putative | 6.74E-05 | -0.55936 |
| PF3D7_0400800 | stevor | 7.76E-05 | -0.55462 |
| PF3D7_0405600 | TMEM33 domain-containing protein, putative | 8.7E-05 | -0.55078 |
| PF3D7_0732400 | rifin | 8.88E-05 | -0.55009 |
| PF3D7_1362000 | cytochrome c oxidase subunit ApiCOX24,<br>putative | 9.32E-05 | -0.54845 |

To understand the transcriptional differences of VSAs, we further analyzed the time course expression of the differentially expressed *var, rif* and *stevor*. In non-adapted parasites, the *var* genes with highest expression differed from each other (PF3D7_1240900 in 3D7-B7, PF3D7_0412400 in 3D7attB, and PF3D7_0420700 in NF54CR) (Table 4), consistent with a previous report (Peters et al., 2007). These three *var* genes belongs to groups B or C var genes and are known to interact with CD36. In contrast, in all three adapted strains, the same *var* gene PF3D7_1200600 (*var2csa*) was the most highly expressed (Table 4 and Figure S1A). *Var2csa* belongs to group E PfEMP1 and does not interact with CD36 (Duffy et al., 2006). Furthermore, 4 *var* genes that belong to group C (Lavstsen et al., 2003) were significantly down-regulated in adapted parasites (Figure S1). Overall, the expression of these differentially expressed *var* genes followed a temporal profile where the transcripts level peaked at 16 hours post infection (hpi), and then dipped to a minimum at 40 hpi (Figures 3A-C, S1B-F), in agreement with previous observations (Bozdech et al., 2003b).

**Table 4.** Most upregulated var in non-adapted and adapted parasites.

| Adaptation | Strain | Gene ID | Group | log2FC |
| --- | --- | --- | --- | --- |
| Non-adapted | 3D7B7 | PF3D7_1240900 | C | 1.16 |
|  | 3D7attB | PF3D7_0412400 | B | 0.80 |
|  | NF54 | PF3D7_0420700 | C | 1.73 |
| Adapted | 3D7B7 AS | PF3D7_1200600 | var2 | 2.33 |
|  | 3D7attB AS | PF3D7_1200600 | var2 | 1.95 |
|  | NF54 AS | PF3D7_1200600 | var2 | 2.33 |

**Figure 3.**
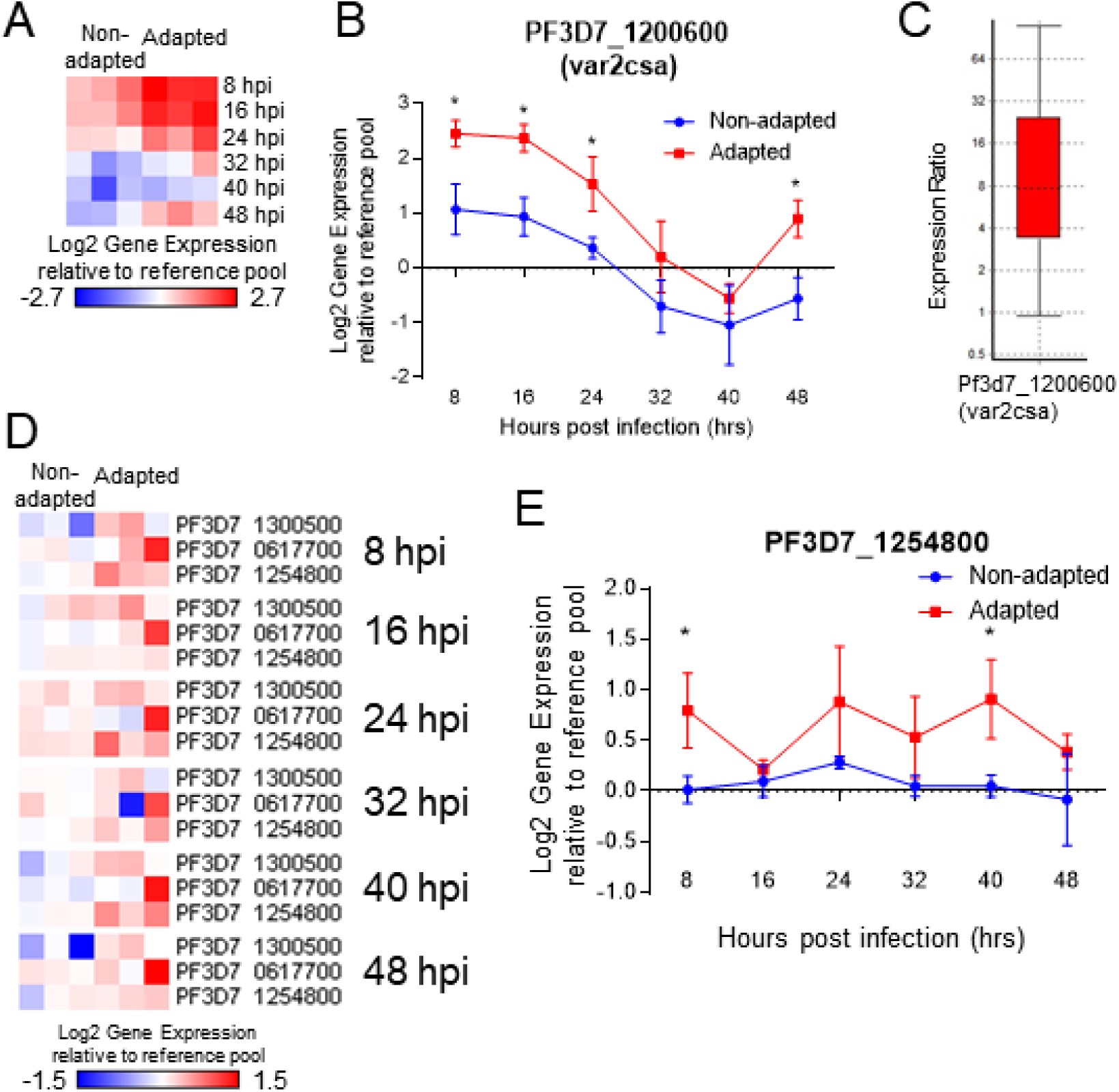
Differentially expressed *var* and *rif* genes between adapted and non-adapted parasites. **(A)** Heatmap of upregulated *var* gene (PF3D7_1200600, *var2csa*). Each column represents a different strain of parasite, while each row represents a different time point of the IDC. **(B)** Temporal gene expression profiles of PF3D7_1200600 across the IDC. Each point represents the mean ± SD. **(C)** qPCR validation of PF3D7_1200600 expression at 16 hours posts post infection (n=5). **(D)** Heatmap of upregulated *rif* genes. Each column represents a different strain of parasite, while each row represents a different *rif* gene. Each cluster represent a different time point of the IDC. (**E**) Temporal gene expression profiles of PF3D7_1254800 across the IDC. Each point represents the mean ± SD. *: p-val<0.05.

Among the *rif* VSA family, 3 type A *rif* were significantly upregulated while 2 were significantly downregulated (Figure S2A). Three type B *rif* were also significantly downregulated (Figure S2B) (Petter, Haeggström et al. 2007). Unlike the *var* genes which exhibited a defined maximum and minimum peak of expression, no such temporal pattern was observed for *rif* genes (Figures 3D-E and S2C-J), consistent with previous observations (Llinas et al., 2006). In addition, 4 *stevor* genes were significantly downregulated in adapted parasites (Figures S3A-E) Like the *rif* genes, the *stevor* genes were continually expressed throughout the IDC with no visible maximal and minimal peak. Taken together, these results show that adaptation is associated with changes in a relatively small number of genes, many of which are the parasite variant surface antigens (VSAs).

### Surface expression of VAS2CSA is required for *in vivo* adaptation through macrophage evasion

To determine the roles of identified DEGs in adaptation of parasites in RBC-NSG mice, we adapted the *in vitro* conditional knockdown TetR-DOZI aptamer system for *in vivo* regulation of *P. falciparum* genes (Ganesan et al., 2016). To validate this approach, 3D7attB AS parasites were transfected with the pMG56 plasmid encoding the firefly luciferase (FLuc) under the translational control of the TetR-DOZI RNA aptamer system, as well as attP sites for Bxb1 integrase-mediated recombination (Figure S4A), to generate 3D7attB AS pMG56 parasites. Human RBCs were infected with 3D7attB AS pMG56 parasites, serially diluted and incubated with luciferin with or without aTc. Luminescent signal was readily detected when 5×10^6^ iRBCs were used (Figure S4B). 3D7attB AS pMG56 parasites were used to infect RBC-NSG mice. When parasitemia in infected mice reached 5%, *in vivo* bioluminescent imaging was performed. Aprroximately 10-fold higher luminescent signal was detected in aTc treated mice (4.7 ± 2.1 ×10^6^ p/s) than in mice without aTc treatment (0.41 ± 0.03 ×10^6^ p/s, unpaired T-test p-value = 0.0251) or uninfected RBC-NSG given aTc (0.44 ± 0.04 ×10^6^ p/s, unpaired T-test p-value = 0.0256) (Figure 4A). These results show that the TetR-DOZI RNA aptamer system can be utilized to regulate the expression of *P. falciparum* proteins in the RBC-NSG mice context.

**Figure 4.**
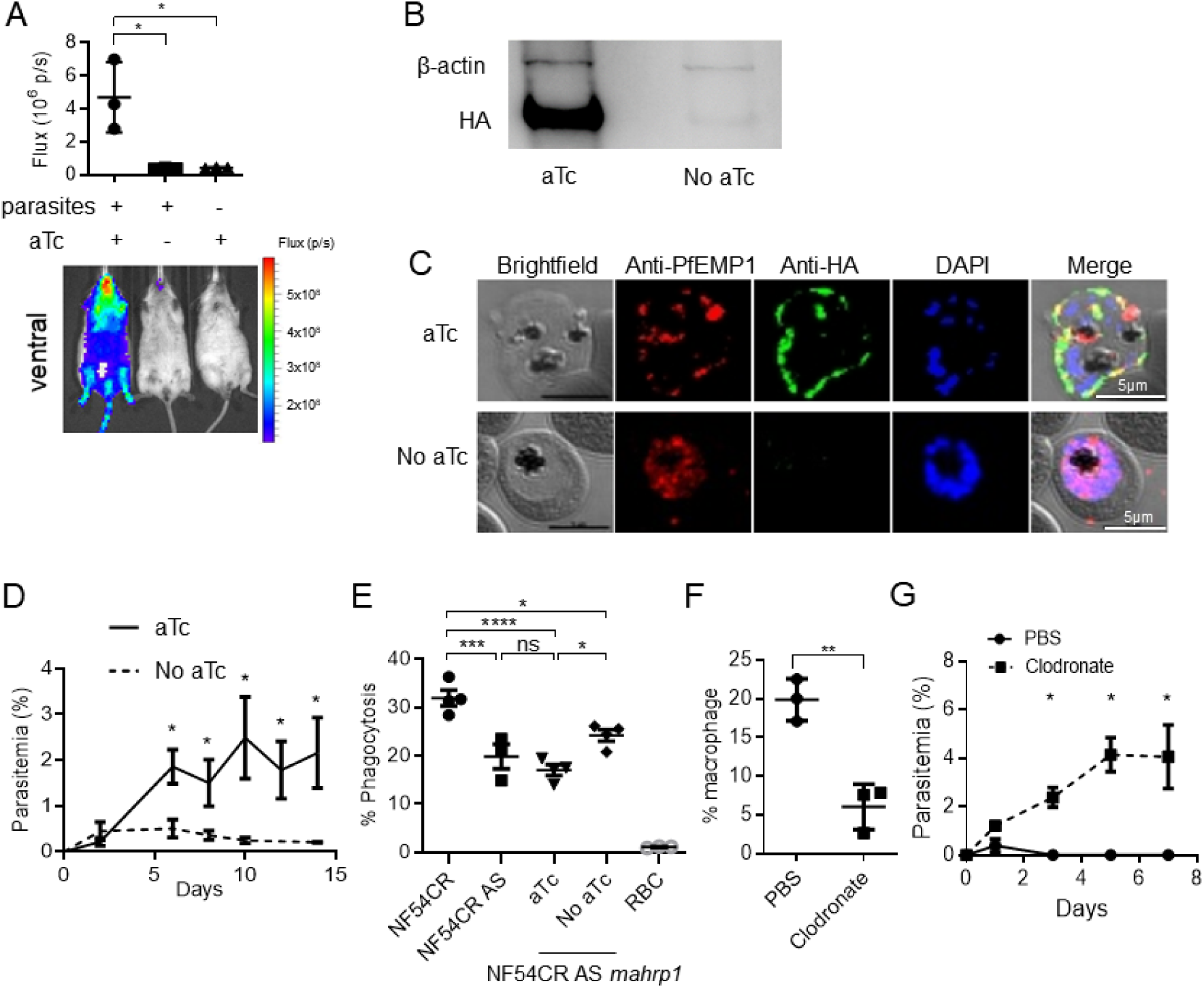
Surface expression of VAR2CSA is required for *in vivo* adaptation. **(A)** RBC-NSG mice were infected with 3D7attB AS pMG56 parasites and were either given aTc or PBS. When parasitemia reached 5%, *in vivo* IVIS imaging to detect luminescence was performed. Data shown are mean ± SEM, n=3. **(B)** Western blot of HA in NF54CR AS *mahrp1* with or without aTc treatment. **(C)** Immunofluorescent assay of NF54CR AS *mahrp1* parasites cultured with or without aTc. Parasites were probed with anti-PfEMP1 (red), anti-HA (green) and DAPI (blue). Scale bar = 5um. **(D)** Parasitemia of RBC-NSG mice infected with NF54CR AS *mahrp1* with (solid) or without (dash) aTc treatment. Parasitemia was assessed by peripheral blood smear with Giemsa staining. Shown are mean parasitemia ± SEM, n=3. **(E)** Comparison of macrophage phagocytosis of NF54CR, NF54CR AS, NF54CR AS *mahrp1* with and without aTc treatment of parasites. Data shown are mean ± SEM, n=3. **(F)** Percentages F4/80^hi^ CD11b^int^ macrophages quantified by flow cytometry in the spleen of RBC-NSG mice given PBS or chlodronate liposone. Data shown are mean ± SEM, n=3. **(G)** Parasitemia of RBC-NSG mice infected with 3D7attB in clodronate-treated mice and control mice treated with PBS-liposome. Parasitemia was assessed by peripheral blood smear with Giemsa staining. Data shown are mean ± SEM, n=3. * p<0.05, ** p<0.01, **, ns=not significant.

To investigate further the biological implication of *var2csa* up-regulation parasite adaptation in RBC-NSG mice, we attempted to target the TetR-DOZI RNA aptamer system into the *var2csa* locus. This approach failed probably because of the high homology among *var* genes at the 3’ ATS segment. Therefore, we used an alternative approach by taking advantage of the fact that members of the PfEMP1 undergo allelic exclusive expression (Voss et al., 2006). We tagged membrane-associated histidine rich protein 1 (MAHRP1, PF3D7_1370300), which is required for PfEMP1 surface expression (Spycher et al., 2008), with the TetR-DOZI system to obtain NF54CR AS *mahrp1* conditional knockdown parasites (Figure S4C). Expression of MAHRP1 was reduced by about 95% in NF54CR AS *mahrp1* in the absence of aTc as compared to the presence of aTc (Figure 4B). Immunofluorescence analyses revealed punctate staining pattern of PfEMP1 close to the surface of the erythrocyte membrane in the presence of aTc (Figure 4C) but a more diffuse staining in the absence of aTc, consistent with an accumulation of PfEMP1 within the parasite plasma membrane. In agreement with previous report (Spycher et al., 2008), no growth defects in these parasites were observed in continuous culture. Mice were infected with NF54CR AS *mahrp1* parasites that had been cultured in the presence of aTc. An initial parasitemia was observed in all infected mice on day 2, which continued to increase in mice that were given aTC (Figure 4D). In contrast, parasitemia in mice without aTc treatment did not increase and eventually reduced to background level. These results suggest that adapted parasites require surface expression of VAR2CSA in order to survive and multiply in RBC-NSG mice.

To investigate the cellular mechanisms involved in VAR2CSA-mediated adaptation, we co-cultured NF54CR, NF54CR AS and NF54CR AS *mahrp1* parasites with human monocyte-derived macrophages (MDM). A phagocytosis rate of 31.9 ± 1.6% (n=4) was observed for non-adapted NF54CR parasites as compared to NF54CR AS parasites (19.8 ± 2.5%, n=3, Tukey’s multiple comparison adjusted p-value = 0.007) (Figure 4E). In the presence of aTc, the phagocytosis rate of NF54CR AS *mahrp1* was 17.0 ± 1.1% (n=4), which was not significantly different from NF54CR AS parasites (Tukey’s multiple comparison adjusted p-value = 0.698). However, in the absence of aTc, the phagocytosis rate was significantly increased to 24.2 ± 1.2% (n=4, Tukey’s multiple comparison adjusted p-value = 0.0221) (Figure 4D). These results suggest that adapted parasites can evade macrophage phagocytosis, probably in part due to selective expression of VAR2CSA that does not bind to CD36 (Serghides et al., 2006).

To further verify the involvement of macrophages in the adaptation of *P. falciparum* parasites, we depleted macrophages in RBC-NSG mice with clodronate liposome (Arnold et al., 2011). After 2 rounds of treatment, the percentage of F4/80^hi^ CD11b^int^ macrophages in the splenic CD45^+^ population was reduced from 19.9 ± 2.7% to 6.1 ± 2.9% (n=3, paired T-test p-value = 0.008) (Figure 4F). The chlodronate-treated mice were infected with non-adapted 3D7 parasites and given chlodronate liposome every two days for the duration of the experiment. In clodronate-treated mice, parasitemia rose to 4.1 ± 2.3% by day 5 while in the untreated mice, no parasites were detectable after day 1 (Figure 4G). Thus, macrophages are a critical immune cell type that parasites must evade during adaptation in RBC-NSG mice.

### Upregulation of immune-modulating RIFINs in adapted parasites leads to decreased NK cell killing but does not affect macrophage phagocytosis

We have previously shown that human NK cells play a critical role in the control of early stages of malaria infection (Chen et al., 2014; Ye et al., 2018). To determine if adapted parasites can resist NK cell-mediated elimination, non-adapted 3D7 or adapted 3D7 AS parasites were co-cultured with NK cells purified from human peripheral blood. While NK cells were able to reduce 3D7 parasitemia by 67.5 ± 3.2% (n=3), this reduction was significantly lowered to 44.3 ± 2.5% for 3D7 AS (n=3, paired T-test p-value = 0.0333) (Figure 5A), demonstrating that adapted parasites were able to evade NK cell killing in vitro.

**Figure 5:**
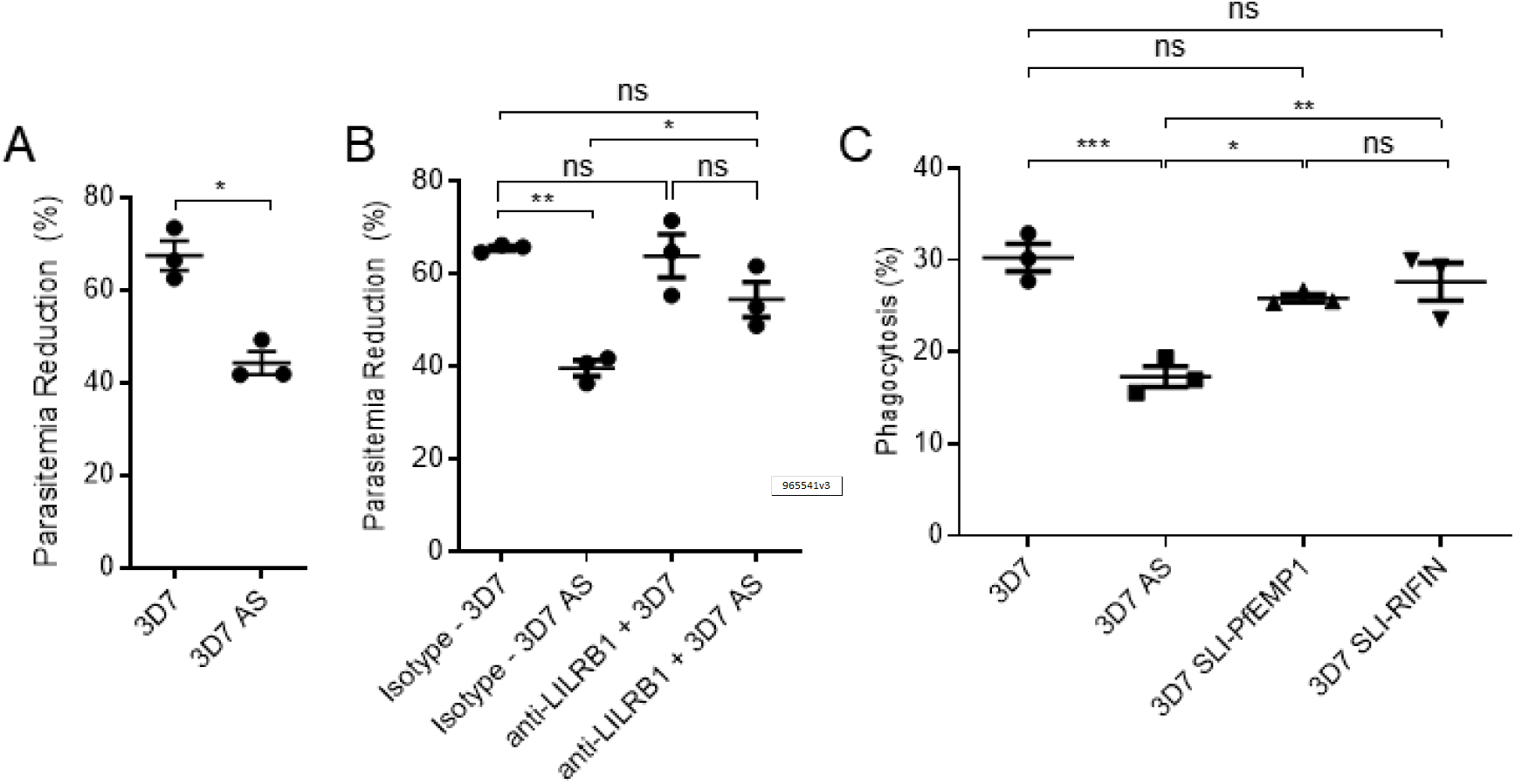
Inhibition of NK cell killing of iRBC through LILRB1 but not in macrophage phagocytosis. **(A)** NK cells were co-cultured with either non-adapted 3D7attB or adapted 3D7attB AS parasites for 96 hrs and parasitemia was quantified by flow cytometry. Parasitemia reduction was calculated as described in materials and methods. **(B)** NK cells were incubated with either an anti-LILRB1 neutralizing antibody or an isotype control and then co-cultured with either 3D7attB or 3D7attB AS parasites for 96 hrs. Parasitemia was quantified by flow cytometry. Data shown sare mean ± SD. **(C)** Macrophage phagocytosis assay of 3D7, 3D7 AS, PfEMP1 PF3D7_0421300-expressing 3D7 (3D7-SLI-PfEMP1) parasites and RIFIN PF3D7_1254800-expressing 3D7 (3D7-SLI-RIFIN). Data shown are mean ± SEM, n=3.

One of the significantly upregulated *rif* gene in our microarray analysis was PF3D7_1254800 (Figure 3D and 3E), which had been shown to bind LILRB1-Fc fusion protein and inhibit NK cell activation (Saito et al., 2017). To determine if parasites evade NK cells through LILRB1, we repeated the co-culture of NK cells with either 3D7 or 3D7 AS parasites in the presence of an anti-LILRB1 neutralizing antibody or an isotype control antibody. As shown in Figure 5B, when NK cells were treated with anti-LILRB1 neutralizing antibody, the parasitemia reduction difference between 3D7 and 3D7 AS was abrogated (63.9 ± 4.7% vs 54.5% ± 3.8, n=3, Tukey’s multiple comparison adjusted p-value = 0.1753). Similarly, when compared to isotype control, the presence of the anti-LILRB1 neutralizing antibody increased parasitemia reduction of 3D7 AS parasites from 39.7 ± 1.7% (n=3) to 54.5 ± 3.8% (n=3, Tukey’s multiple comparison adjusted p-value = 0.0342).

We have recently utilized a selection-linked integration (SLI) approach (Birnbaum et al., 2017) to select for non-adapted 3D7 parasites that expressed PF3D7_1254800 (3D7-SLI-RIFIN) and verified its role in the modulation NK cell mediated killing (Omelianczyk et al., 2020). To evaluate whether PF3D7_1254800 also plays a role in reducing phagocytosis by macrophages we utilized the 3D7-SLI-RIFIN along with a parasite line that expressed the SLI tagged *var* gene (PF3D7_0421300) to produce 3D7-SLI-PfEMP1 parasites. Macrophage phagocytosis levels were similar to non-adapted 3D7 parasites (30.3 ± 2.6%, n=3) for both 3D7-SLI-RIFIN (27.6 ± 3.5%, n=3) as well as 3D7-SLI-PfEMP1 (25.9 ± 0.7%, n=3) as compared to 3D7 AS (17.3 ± 1.9%, n=3) which showed significantly reduced phagocytosis levels compared to 3D7 (Tukey’s multiple comparison adjusted p-value = 0.0008), 3D7-SLI-RIFIN (Tukey’s multiple comparison adjusted p-value = 0.0036) and 3D7-SLI-PfEMP1 (Tukey’s multiple comparison adjusted p-value = 0.0109) (Figure 5C). These results show that the ability of adapted parasites to evade NK cell killing is associated with upregulation of the RIFIN PF3D7_1254800, which inhibits NK cell activation through LILRB1. However, upregulation of RIFIN PF3D7_1254800 does not reduce macrophage phagocytosis of non-adapted parasites.

## Discussion

Over the course of co-evolution with its human host, *P. falciparum* has developed a repertoire of mechanisms to evade the host immune clearance. One of such mechanisms is the switching of its surface-exposed VSA, especially VSAs belonging to the *var* family. Expression of *var* in freshly isolated parasites from patients was observed to be highly coordinated, with a single dominant var gene being expressed at any time. However, such coordination is lost during continuous *in vitro* culture, resulting in a random non-coordinated expression of *var* genes (Bachmann et al., 2011). The overall expression of *var* genes are also downregulated (Peters, Fowler et al. 2007). Here, we report that *in vivo* adaptation of parasites resulted in unique phenotypic and transcriptomic changes in the malaria parasite. Notably, there was an almost four-fold upregulation of the *var2csa* in adapted parasites, coupled with downregulation of four CD36-binding Group C *var* genes (Janes et al., 2011).

*Var2csa* is one of three *vars* that are conserved across the different *P. falciparum* strains (Rowe, Kyes et al. 2002). VAR2CSA is the only member of the group E PfEMP1 (Salanti et al., 2003) and binds chondroitin sulphate A (CSA) but not CD36 (Salanti et al., 2003). Studies have shown that parasites isolated from patients with severe malaria express high levels of group A PfEMP1 that do not bind CD36 (Bernabeu et al., 2016; Rottmann et al., 2006), suggesting a role of avoiding recognition by CD36 is a mechanism of parasite evasion of macrophage phagocytosis. We have now demonstrated this mechanism of action beyond previous correlation. Parasites that successfully adapted to grow in RBC-NSG mice expressed VAR2CSA and VAR2CSA-expressing adapted parasites were resistant to macrophage phagocytosis *in vitro*. When VAR2CSA expression was inhibited, adapted parasites were efficiently phagocytosed by macrophages *in vitro* and did not induce patent infection in RBC-NSG mice. Conversely, when macrophages were depleted, non-adapted parasites could immediately flourish in RBC-NSG mice. Together, these results demonstrate that expression of PfEMP1, such as VAR2CSA, that does not bind to CD36, is a mechanism by which parasites evade macrophages in order to establish robust infection *in vivo*.

RIFINs also show differential expression during the adaptation process. The type A RIFINs are localized to the iRBC membrane and might be surface exposed, while type B RIFINs remain inside the infected RBC (Bachmann et al., 2015; Petter et al., 2007). We found that all upregulated *rif* genes are A-type, while three out of five downregulated *rif* are B-type. Anti-RIFIN antibodies have been shown to impact disease severity (del Pilar Quintana et al., 2018), suggesting a role of RIFIN in malaria pathology. In particular, a subset of RIFIN have been shown to inhibit NK cell activation through the inhibitory receptor LILRB1 receptor (Saito et al., 2017). We show that a consistently upregulated RIFIN in adapted parasites is PF3D7_1254800. This RIFIN mediates parasite escape from NK cell killing by inhibiting LILRB1 as blocking LILRB1 by a neutralizing antibody prevents the escape. Thus, our findings identify a specific RIFIN that mediates escape from NK cells during parasite adaptation.

It is notable that despite the absence of NK cells in the RBC-NSG model, we were able to detect the specific upregulation of *rif* genes that are involved in evasion of NK cells. One possibility is that there is a so far unknown coordination of *var* and *rif* expression that provides the parasites with the ability to escape the host innate immune response. Such a coordinated expression could be particularly important during the early stage of parasite infection when relatively small numbers of parasites are released from the liver schizont and are particularly at risk from the innate clearance. Such a unique survival strategy has also been proposed for liver stage merozoites in the rodent malaria parasite *P. yoelii*, that express a unique member of an invasion ligand to facilitate the initial invasion of RBCs (Carlton et al., 2002). Although how this transcriptional co-regulation is achieved is at this point not known, it is possible that an epigenetic immune-evasion program is activated during the adaptation process as members of the *var, rif* and *stevor* share common transcriptional activators (Howitt et al., 2009) (Karmodiya et al., 2015). Such an epigenetic imprinting could also allow for adapted parasites to maintain their phenotype even after more than 20 cycles of *in vitro* culture.

In conclusion, we show that significant changes in transcription occur during parasite adaptation in RBC-NSG mice. These changes are not random but specifically happen to genes involved in immune evasion. In particular, a single *var* known to be involved in repressing macrophage phagocytosis and a small set of *rif* important to escape NK cell killing were repeatedly upregulated in multiple adapted parasites, suggesting immune evasion during adaptation. As escape from clearance by the innate immune system is particularly important early on during the establishment of the blood stage infection when parasite numbers are still relatively small, identfication of specific *var* and *rif* that are upregulated during parasite adaptation provide targets for developing more effective malaria vaccines.

## Materials and Methods

### *P. falciparum* Strains and culturing

*P. falciparum* blood-stage parasites were cultured in 2.5% hematocrit human RBC in malaria culture media (MCM) comprising of 10.43g RPMI 1640 powder (Gibco), 25ml 1M HEPES (Gibco), 2g NaHCO_3_ (Sigmaaldrich), 5g Albumax (Gibco), 0.05g hypoxanthine (Sigma-aldrich) and 25mg gentamicin (Gibco) in 1L milli-Q water. Cultures were incubated in Heracell 150 incubator (Thermo Scientific) at 37 in 5% CO2, 3% O2 and 92% N2.

NF54CR (NF54^Cas9+T7 Polymerase^) (Nasamu et al., 2019) parasites were kindly shared with us by Prof. Jacquin Niles. Other parasite lines used were obtained from BEI Resources (Table 1).

### Primary Cells

Whole blood donated by healthy non-malarial immune adult volunteers at the National University Hospital of Singapore Blood Donation Center. Informed consent was obtained from all donors in accordance with approved protocol and guidelines. Project ethics and approval were obtained from the Institutional Review Board of National University of Singapore (NUS-IRB 10-285). Whole venous blood collected in Citrate-Phosphate Dextrose-Adenine-1 (CPDA-1,JMS) and PBMC were isolated from whole blood over Ficoll-Paque PLUS (GE Healthcare) density gradient. Pelleted RBC were washed twice in RPMI 1640 (Sigma-Aldrich) and stored 1:1 in MCM. Remaining RBCs within the PBMC fraction were lysed with ACK lysis buffer (Life Technologies) and purified PBMC were washed twice with RPMI 1640. PBMCs were counted and cryopreserved at a concentration of 1×10^8^ cells/ml in 1:1 RPMI and PBMC freezing medium of 85% fetal bovine serum (Gibco) and 15% dimethyl sulfoxide (Sigma-Aldrich) in liquid nitrogen.

### Mice

6-week old NOD/SCID IL2γnull mice were approved by the institutional animal care and use committee (IACUC) of National University of Singapore (NUS), Agency of Science, Technology and Research (A*STAR) and Massachusetts Institute of Technology (MIT).

### Plasmids

Plasmids pMG56 (Ganesan et al., 2016) and pSN054 were kindly shared by Prof Jacquin Niles for the generation of 3D7attb pMG56 and NF54CR AS *mahrp1* parasites respectively.

### *In vivo* infection

RBC-NSG mice were generated as described previously (Zenonos et al., 2015). Briefly, 6-week-old NSG mice were injected daily with RBC mixture (50% human RBC, 25% RPMI, 25% human AB serum) intraperitoneally. Human RBC reconstitution was assessed every other day by determining CD235ab levels via flow cytometry. Mice with reconstitution levels above 20% were infected with 1 × 10^7^ mixed stage *P. falciparum* intravenously. Peripheral blood parasitemia was determined via Giemsa Staining. Upon detection of parasitemia, mice were bled and adapted parasites were recovered.

For macrophage depletion, RBC-NSG mice were treated with 25ul of Clodrosome® Liposomal Clodronate (5mg/ml) (Encapsula NanoSciences LLC) intraperitoneally at Day -6, Day-4 and Day -2 before infection with *P*.*falciparum* at Day 0.

*In vivo* imaging was performed using an IVIS® Spectrum (PerkinElmer) on mice injected with RediJect D-Luciferin Bioluminescent Substrate (PerkinElmer) at 100ul via intraperitoneal injection. Data was analyzed on LivingImage 3.2 (PerkinElmer).

### Whole genome sequencing

*P. falciparum* genomic DNA was isolated using NucleoSpin® Blood Columns (Macherey-Nagel) as per manufacturer’s protocol. Whole genome next generation sequencing of adapted and non-adapted *P. falciparum* was performed at Singapore Centre for Environmental Life Sciences Engineering Sequencing Core using Illumina Miseq Run V3. FASTQ files of sequencing reads were aligned to the *P. falciparum* 3D7 reference genome available, at PlasmoDB, using bowtie2 v2.3.2. SAM file generated from the alignment is converted to BAM files using samtools v1.3. SNP variant calling on the BAM files was done using freebayes v1.0.1 and SNP filtering based on QUAL using vcffilter.

### Transcriptional microarray analysis

Microarray analysis was performed as described previously (Bozdech et al., 2003b). Briefly, *P. falciparum* RNA was isolated using TRIzol reagent (Invitrogen) as per manufacturer’s protocol. RNA integrity was determined using 2100 Bioanalyzer with RNA 6000 Nano chips (Agilent). cDNA was synthesized using a combination of SMARTer PCR cDNA Synthesis Kit (Takara) and SuperScript II Reverse Transcriptase (ThermoFisher). cDNA was then amplified in the presence of 0.225mM amino-allyl-dUTP (Biotium) using Taq DNA polymerase (New England Biolabs). PCR products were purified with MinElute PCR purification kit (Qiagen) according to manufacturer’s protocol and eluted in 16 µl of elution buffer. The reference pool was created by adding equal amounts of RNA from each timepoint from the non-adapted strain.

4 µg of amplified DNA samples were labelled with Cy3 or Cy5 (GE Healthcare). Experimental samples were labeled with Cy5 while reference pools were labeled with Cy3. Cy5-labeled and Cy3-labeled reference pool were mixed and hybridized on post-processed microarray chips using the Agilent hybridization system (Agilent), and scanned using PowerScannerTM (Teacan). Scanned images were analyzed using GenePix Pro 6.0 (Axon Instruments). Microarray data was LOESS normalized and filtered for signal intensity over the background noise using R package LIMMA (Ritchie et al., 2015). Differentially expressed genes were determined using Significance Analysis of Microarray (SAM) (Tusher et al., 2001).

### Peripheral blood mononuclear cell (PBMC) purification

PBMCs were isolated using Ficoll-Paque PLUS (GE Healthcare) as per manufacturer protocol. NK cells and monocytes were isolated from purified PBMCs using EasySep™ Human NK Cell Isolation Kit and EasySep™ Human Monocyte Isolation Kit (STEMCELL) respectively.

### Phagocytosis assay

Monocyte-derived macrophages were obtained by culturing purified monocytes in RPMI 10% FBS for 7 days. Non-adherent cells are removed. Phagocytosis assay was performed by incubating DAPI-treated trophozoites at a ratio of 5 iRBCs to 1 macrophage for 90 mins. Thereafter, the co-culture was washed in PBS to remove excess iRBCs. Macrophages were detached using Accutase (StemCell Technologies), stained with anti-human CD14 (Clone 63D3; Biolegend) and quantified using Attune NxT (Life Technologies). Percentage phagocytosis is calculated as the percentage of DAPI-positive macrophages.

### Primary NK cell parasitemia reduction co-culture

NK cell killing assay was performed as previously described (Chen et al., 2014). Briefly, primary NK cells were co-cultured with trophozoite at a parasitemia of 0.5% and at a ratio of 1 iRBCs to 10 NK cells for 96 hrs. Quantification of parasitemia was done by flow cytometry. Parasitemia reduction was calculated as 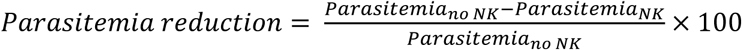. Neutralization of LILRB1 was performed with anti-LILRB1 antibody (500ng/ml, R&D Systems, MAB20172).

### Immunofluorescence assay (IFA)

Smears of late stage parasite cultures were made on glass slides and methanol fixed on ice for 15 mins and air dried. Smears were blocked in 3% bovine serum albumin (BSA; Sigma-aldrich). Slides were incubated with primary rat anti-HA (Roche) at 1:100 and primary mouse anti-PfEMP1 ATS at 1:500 overnight. After three rounds of washing, slides were incubated with secondary goat anti-rat IgG (H+L) Alexa Fluor 488 (1:500; Invitrogen) and goat anti-mouse IgG (H+L) Alexa Fluor 647 (1:500; Invitrogen) with Hoechst 33342 (1:1000) for 1 hour at room temperature. The slides were then mounted in VECTASHIELD® Antifade mounting media (Vector Laboratories), imaged on LSM710 confocal microscope (Carl Zeiss) and analyzed on ZEN 2 (Carl Zeiss).

### Western Blot

Late trophozoite parasites cultured with or without aTc were isolated using a 65% Percoll gradient. Recovered parasites were resuspended in Laemmli Sample Buffer (Bio-rad) and β-mercaptoethanol (Sigma-aldrich) and loaded on a 10% Mini-PROTEAN® TGX™ Precast Protein Gel. Samples were transferred to 0.2 μm Polyvinylidene difluoride (PVDF) membrane using Trans-Blot Turbo Transfer System (Bio-rad). The membrane was blocked in 5% skim milk in 0.1% PBS-Tween (PBS-T) for 1 hour at room temperature, and incubated overnight with rat anti-HA tag (Roche) and mouse anti-Actin (Invitrogen) at 1:3000 in 2% Bovine Serum Albumin (BSA). The blot was then washed 3 times in PBS-T and probed with goat anti-rat HRP (Biolegend) and goat anti-mouse HRP (Biolegend) at 1:10,000 in 2% BSA PBS-T for 1 hour at room temperature. The blot was imaged on ChemiDoc MP (Bio-rad) in Clarity Max Western ECL Substrate (Bio-rad). Western blot analysis was done using Image Lab v6.0 (Bio-rad).

### Quantification and Statistical Analysis

Data are presented as the mean and standard error of mean (SEM). Differences between paired samples were analyzed using a paired t-test, while unpaired samples were analyzed with Student’s t-test. Multiple comparison tests was performed using Tukey’s multiple comparison test. A p-value of < 0.05 was considered statistically significant. All calculations were performed using the GraphPad 6.01 software package.

## Acknowledgments

We thank Lan Hiong Wong for technical assistance, Farzad Olfat for administrative support and Neslihan Kaya for bioinformatics analysis. This work was supported by the National Research Foundation of Singapore through the Singapore–MIT Alliance for Research and Technology’s (SMART) Interdisciplinary Research Groups in Infectious Disease and Antimicrobial Resistance Research Program and the Singapore Ministry of Health’s National Medical Research Council under its Cooperative Basic Research Grant (NMRC/CBRG/0040/2013) and Open Fund Individual Research Grant (NMRC/OFIRG/0058/2017). M.C. and W.Y. were supported by the SMART graduate fellowship while R.I.O. was supported by a SINGA graduate fellowship.

## Author Contributions

Conceptualization, M.C., W.Y., P.P. and J.C.; Methodology, M.C., W.Y., C.F.P., R.H., J.C.N.; Investigation, M.C. and W.Y.; Writing – Original Draft, M.C., W.Y.; Writing – Review & Editing, M.C., W.Y., P.P. and J.C.; Funding Acquisition, P.P. and J.C.; Resources, Q.C., J.C.N., R.I.O., P.P and J.C.; Supervision, P.P and J.C.

## Supplemental Figures

**Figure S1:**
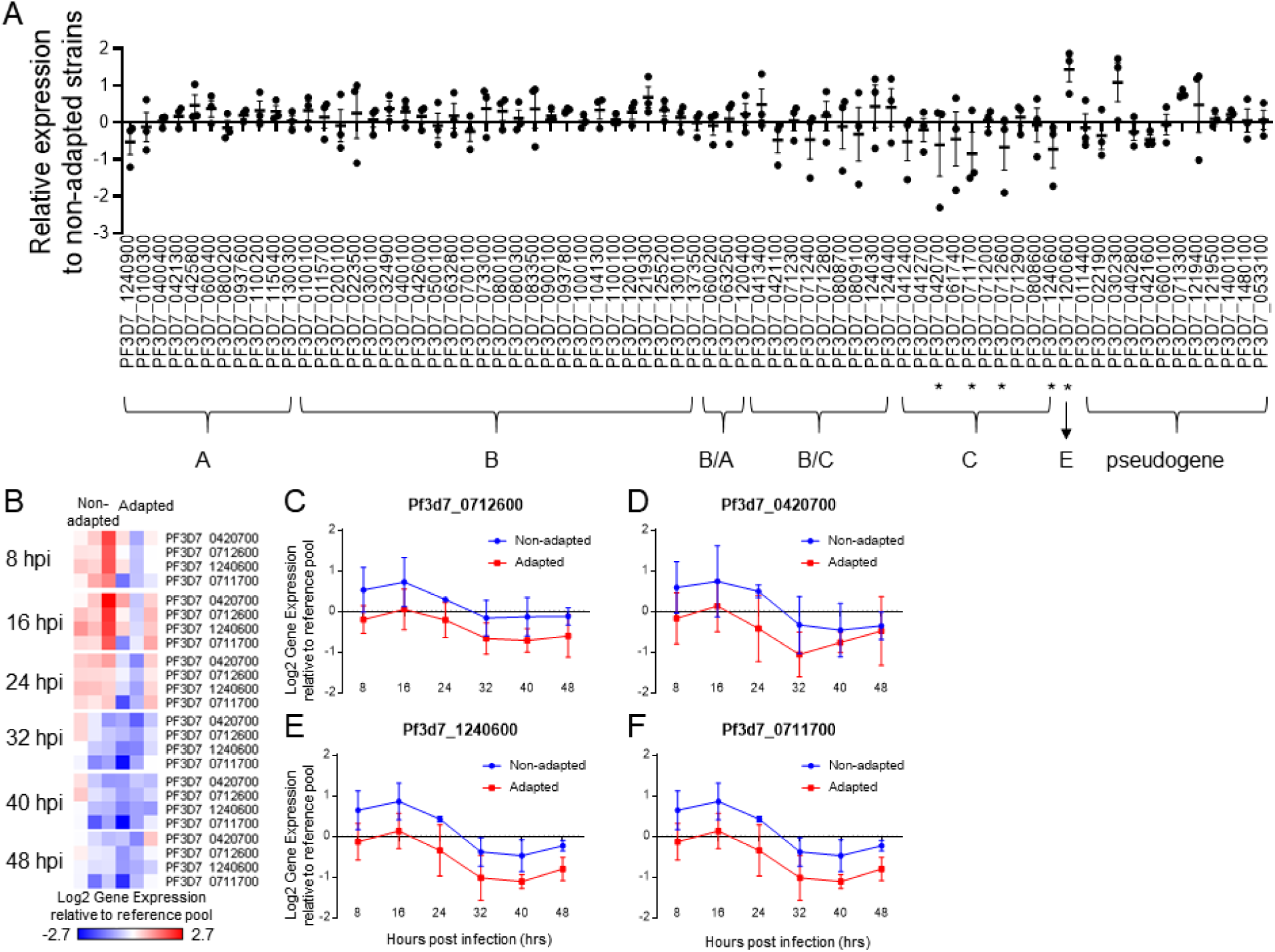
Differential expression of *var* genes between adapted and non-adapted parasites. **(A)** Relative expression of individual *var* genes in adapted parasites compared to non-adapted parasites. *Var* genes are grouped according to the classification proposed by Lavstsen *et al* (Lavstsen et al., 2003). Each point represents a different pair of adapted and non-adapted parasites. *: differentially expressed gene. **(B)** Heatmap of the downregulated *var* genes. Each column represents a different strain of parasite, while each row represents a different *var* gene. Each cluster represent a different time point of the IDC. **(C-F)** Temporal gene expression profiles of the indicated *var* gene across the IDC. Each point represents the mean±SD.

**Figure S2:**
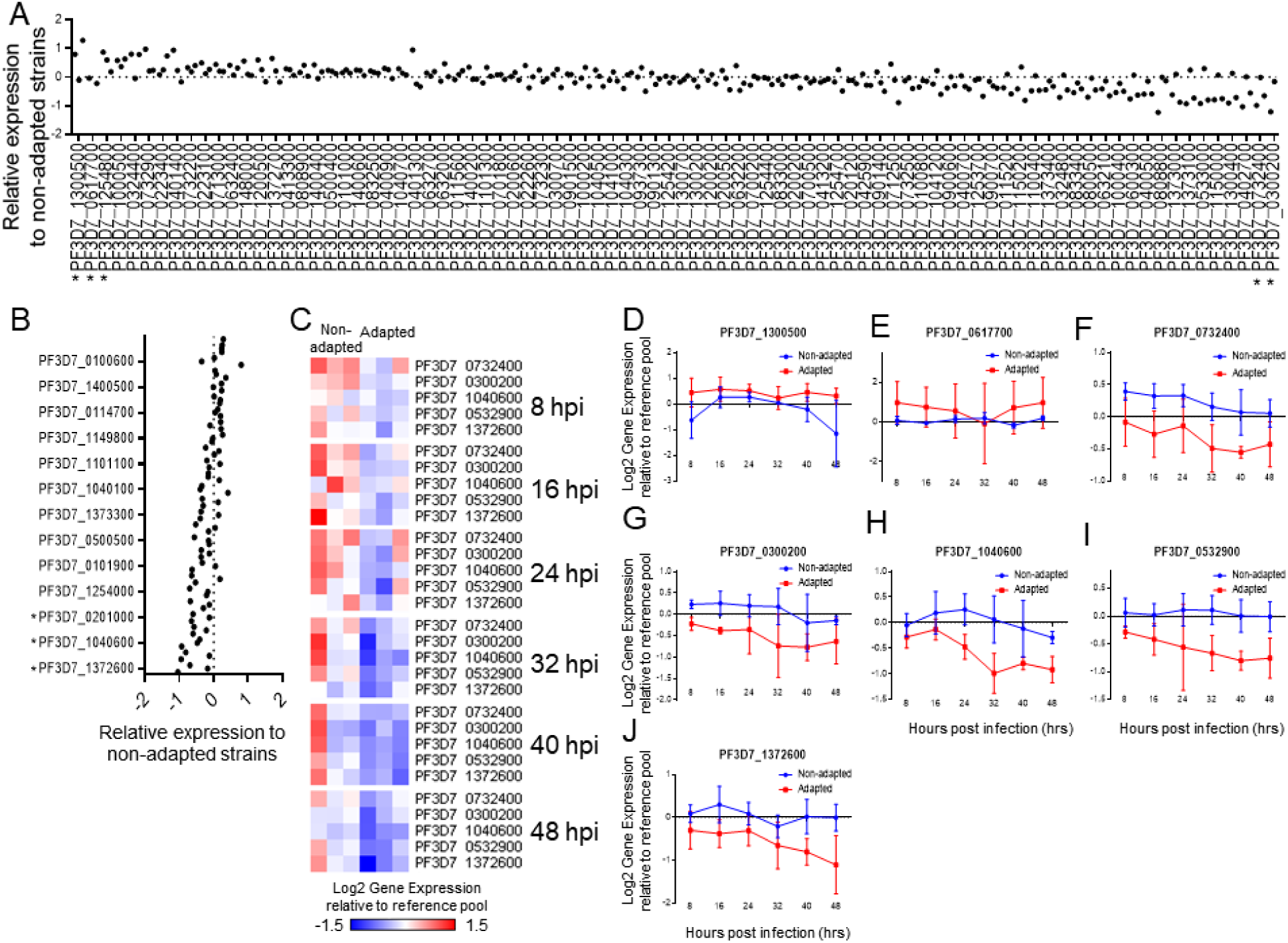
Differential expression of *rif* genes between adapted and non-adapted parasites. **(A-B)** Relative expression of type A **(A)** and type B **(B)** *rif* genes in adapted parasites compared to non-adapted parasites. Each point represents a different pair of adapted and non-adapted parasites. *: differentially expressed gene. **(C)** Heatmap of downregulated *rif* genes. Each column represents a different strain of parasite, while each row represents a different *rif* gene. Each cluster represent a different time point of the IDC. **(D-J)** Temporal gene expression profiles of the indicated *rif* gene across the IDC. Each point represents the mean ± SD.

**Figure S3.**
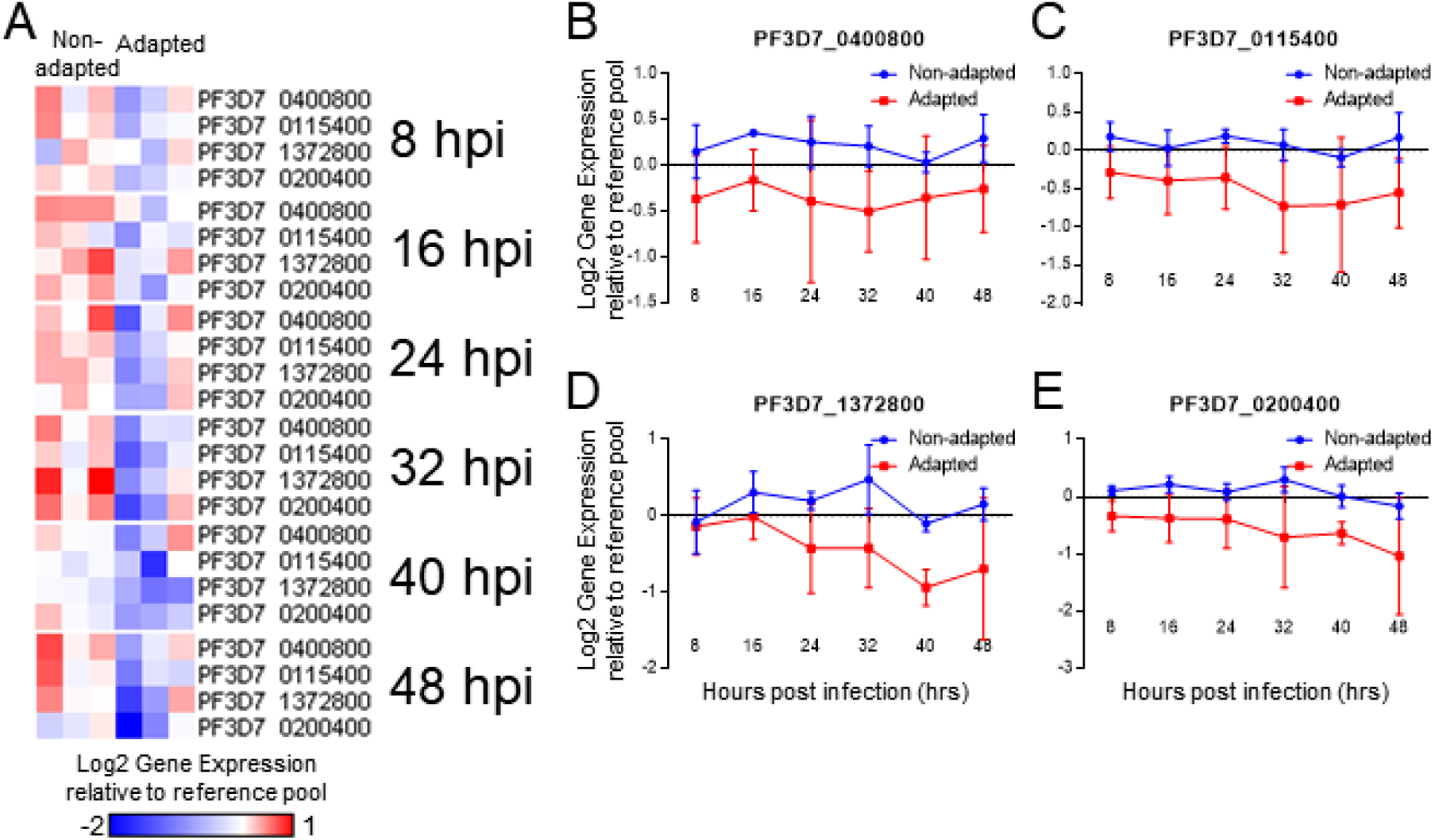
Differentially expressed *stevor* genes between adapted and non-adapted parasites. **(A)** Heatmap of downregulated *stevor* genes. Each column represents a different strain of parasite, while each row represents a different *rif* gene. Each cluster represent a different time point of the IDC. (**B-E**) Temporal gene expression profiles of the indicated gene across the IDC. Each point represent the mean ± SD.

**Figure S4.**
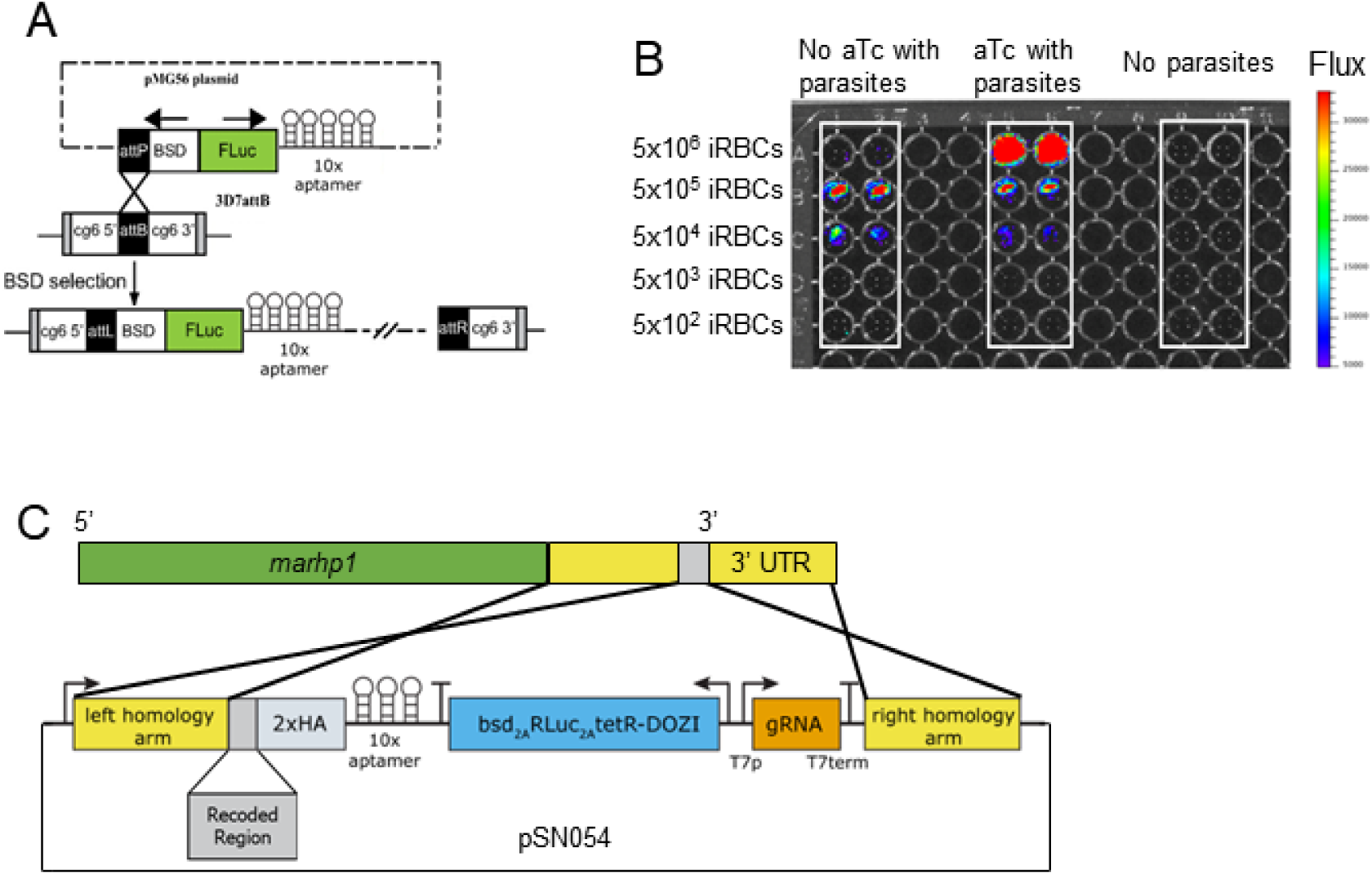
Schematics of plasmids used. **(A)** Schematic of pMG56 plasmid carrying the firefly luciferase (FLuc) gene under the translational control of TetR-DOZI RNA aptamer system as previously described (Ganesan et al., 2016). AttP containing pMG56 was integrated into attB containing 3D7attB AS *P. falciparum* parasites at the dispensable cg6 gene locus to generate 3D7attB AS pMG56 parasites. **(B)** 3D7attB AS pMG56 parasites were serially diluted in a 96 well plate with luciferin and either in the presence or absence of aTc. Bioluminescence was analyzed using IVIS. **(C)** Schematic of plasmid pSN054 used to introduce a 2xHA tag as well as the 10x aptamer array to the 3’ end of the *mahrp1* gene as well as components for translation regulation by the TetR-DOZI system.

## References

Abdel-Latif, M.S., Khattab, A., Lindenthal, C., Kremsner, P.G., and Klinkert, M.Q. (2002). Recognition of variant Rifin antigens by human antibodies induced during natural Plasmodium falciparum infections. Infection and immunity 70, 7013–7021.

Adjalley, S.H., Johnston, G.L., Li, T., Eastman, R.T., Ekland, E.H., Eappen, A.G., Richman, A., Sim, B.K.L., Lee, M.C., and Hoffman, S.L. (2011). Quantitative assessment of Plasmodium falciparum sexual development reveals potent transmission-blocking activity by methylene blue. Proceedings of the National Academy of Sciences 108, E1214–E1223.

Angulo-Barturen, I., Jiménez-Díaz, M.B., Mulet, T., Rullas, J., Herreros, E., Ferrer, S., Jiménez, E., Mendoza, A., Regadera, J., Rosenthal, P.J., et al. (2008). A Murine Model of Falciparum-Malaria by In Vivo Selection of Competent Strains in Non-Myelodepleted Mice Engrafted with Human Erythrocytes. PLoS ONE 3, e2252.

Arnold, L., Tyagi, R.K., Meija, P., Swetman, C., Gleeson, J., Perignon, J.L., and Druilhe, P. (2011). Further improvements of the P. falciparum humanized mouse model. PLoS One 6, e18045.

Ashburner, M., Ball, C.A., Blake, J.A., Botstein, D., Butler, H., Cherry, J.M., Davis, A.P., Dolinski, K., Dwight, S.S., and Eppig, J.T. (2000). Gene ontology: tool for the unification of biology. Nature genetics 25, 25.

Bachmann, A., Predehl, S., May, J., Harder, S., Burchard, G.D., Gilberger, T.W., Tannich, E., and Bruchhaus, I. (2011). Highly co-ordinated var gene expression and switching in clinical Plasmodium falciparum isolates from non-immune malaria patients. Cellular microbiology 13, 1397–1409.

Bachmann, A., Scholz, J.A.M., Janßen, M., Klinkert, M.-Q., Tannich, E., Bruchhaus, I., and Petter, M.J.M.j. (2015). A comparative study of the localization and membrane topology of members of the RIFIN, STEVOR and Pf MC-2TM protein families in Plasmodium falciparum-infected erythrocytes. 14, 274.

Bernabeu, M., Danziger, S.A., Avril, M., Vaz, M., Babar, P.H., Brazier, A.J., Herricks, T., Maki, J.N., Pereira, L., Mascarenhas, A., et al. (2016). Severe adult malaria is associated with specific PfEMP1 adhesion types and high parasite biomass. Proceedings of the National Academy of Sciences 113, E3270.

Birnbaum, J., Flemming, S., Reichard, N., Soares, A.B., Mesen-Ramirez, P., Jonscher, E., Bergmann, B., and Spielmann, T. (2017). A genetic system to study Plasmodium falciparum protein function. Nature methods 14, 450–456.

Bozdech, Z., Llinás, M., Pulliam, B.L., Wong, E.D., Zhu, J., and DeRisi, J.L. (2003a). The Transcriptome of the Intraerythrocytic Developmental Cycle of Plasmodium falciparum. PLOS Biology 1, e5.

Bozdech, Z., Zhu, J., Joachimiak, M.P., Cohen, F.E., Pulliam, B., and DeRisi, J.L. (2003b). Expression profiling of the schizont and trophozoite stages of Plasmodium falciparum with a long-oligonucleotide microarray. Genome biology 4, R9.

Bull, J.J., Molineux, I.J., and Rice, W.R. (1991). SELECTION OF BENEVOLENCE IN A HOST-PARASITE SYSTEM. Evolution; international journal of organic evolution 45, 875–882.

Carlton, J.M., Angiuoli, S.V., Suh, B.B., Kooij, T.W., Pertea, M., Silva, J.C., Ermolaeva, M.D., Allen, J.E., Selengut, J.D., Koo, H.L., et al. (2002). Genome sequence and comparative analysis of the model rodent malaria parasite Plasmodium yoelii yoelii. Nature 419, 512–519.

Chan, J.A., Howell, K.B., Reiling, L., Ataide, R., Mackintosh, C.L., Fowkes, F.J., Petter, M., Chesson, J.M., Langer, C., Warimwe, G.M., et al. (2012). Targets of antibodies against Plasmodium falciparum-infected erythrocytes in malaria immunity. The Journal of clinical investigation 122, 3227–3238.

Chen, Q., Amaladoss, A., Ye, W., Liu, M., Dummler, S., Kong, F., Wong, L.H., Loo, H.L., Loh, E., Tan, S.Q., et al. (2014). Human natural killer cells control Plasmodium falciparum infection by eliminating infected red blood cells. Proceedings of the National Academy of Sciences of the United States of America 111, 1479–1484.

Consortium, G.O. (2018). The gene ontology resource: 20 years and still GOing strong. Nucleic acids research 47, D330–D338.

Day, K.P., Karamalis, F., Thompson, J., Barnes, D.A., Peterson, C., Brown, H., Brown, G.V., and Kemp, D.J. (1993). Genes necessary for expression of a virulence determinant and for transmission of Plasmodium falciparum are located on a 0.3-megabase region of chromosome 9. Proceedings of the National Academy of Sciences of the United States of America 90, 8292–8296.

del Pilar Quintana, M., Ch’ng, J.-H., Moll, K., Zandian, A., Nilsson, P., Idris, Z.M., Saiwaew, S., Qundos, U., and Wahlgren, M.J.S.r. (2018). Antibodies in children with malaria to PfEMP1, RIFIN and SURFIN expressed at the Plasmodium falciparum parasitized red blood cell surface. 8, 1–14.

Dodoo, D., Staalsoe, T., Giha, H., Kurtzhals, J.A., Akanmori, B.D., Koram, K., Dunyo, S., Nkrumah, F.K., Hviid, L., and Theander, T.G. (2001). Antibodies to variant antigens on the surfaces of infected erythrocytes are associated with protection from malaria in Ghanaian children. Infection and immunity 69, 3713–3718.

Duffy, M.F., Maier, A.G., Byrne, T.J., Marty, A.J., Elliott, S.R., O’Neill, M.T., Payne, P.D., Rogerson, S.J., Cowman, A.F., Crabb, B.S., et al. (2006). VAR2CSA is the principal ligand for chondroitin sulfate A in two allogeneic isolates of Plasmodium falciparum. Molecular and biochemical parasitology 148, 117–124.

Fenton, B., Clark, J.T., Wilson, C.F., McBride, J.S., and Walliker, D. (1989). Polymorphism of a 35-48 kDa Plasmodium falciparum merozoite surface antigen. Molecular and biochemical parasitology 34, 79–86.

Fernandez, V., Hommel, M., Chen, Q., Hagblom, P., and Wahlgren, M. (1999). Small, clonally variant antigens expressed on the surface of the Plasmodium falciparum-infected erythrocyte are encoded by the rif gene family and are the target of human immune responses. The Journal of experimental medicine 190, 1393–1404.

Ganesan, S.M., Falla, A., Goldfless, S.J., Nasamu, A.S., and Niles, J.C. (2016). Synthetic RNA–protein modules integrated with native translation mechanisms to control gene expression in malaria parasites. Nat Commun 7, 10727.

Gomes, P.S., Bhardwaj, J., Rivera-Correa, J., Freire-De-Lima, C.G., and Morrot, A. (2016). Immune Escape Strategies of Malaria Parasites. Front Microbiol 7, 1617-1617.

Gross, M. (2019). Fresh efforts needed against malaria. Current Biology 29, R301–R303.

Howitt, C.A., Wilinski, D., Llinás, M., Templeton, T.J., Dzikowski, R., and Deitsch, K.W. (2009). Clonally variant gene families in Plasmodium falciparum share a common activation factor. Molecular microbiology 73, 1171–1185.

Janes, J.H., Wang, C.P., Levin-Edens, E., Vigan-Womas, I., Guillotte, M., Melcher, M., Mercereau-Puijalon, O., and Smith, J.D. (2011). Investigating the Host Binding Signature on the Plasmodium falciparum PfEMP1 Protein Family. PLOS Pathogens 7, e1002032.

Karmodiya, K., Pradhan, S.J., Joshi, B., Jangid, R., Reddy, P.C., and Galande, S. (2015). A comprehensive epigenome map of Plasmodium falciparum reveals unique mechanisms of transcriptional regulation and identifies H3K36me2 as a global mark of gene suppression. Epigenetics & Chromatin 8, 32.

Lai, S.M., Sheng, J., Gupta, P., Renia, L., Duan, K., Zolezzi, F., Karjalainen, K., Newell, E.W., and Ruedl, C. (2018). Organ-specific fate, recruitment, and refilling dynamics of tissue-resident macrophages during blood-stage malaria. Cell reports 25, 3099-3109. e3093.

Langreth, S.G., and Peterson, E. (1985). Pathogenicity, stability, and immunogenicity of a knobless clone of Plasmodium falciparum in Colombian owl monkeys. Infection and immunity 47, 760.

Lavstsen, T., Salanti, A., Jensen, A.T.R., Arnot, D.E., and Theander, T.G. (2003). Sub-grouping of Plasmodium falciparum 3D7 var genes based on sequence analysis of coding and non-coding regions. Malaria journal 2, 27–27.

Llinas, M., Bozdech, Z., Wong, E.D., Adai, A.T., and DeRisi, J.L. (2006). Comparative whole genome transcriptome analysis of three Plasmodium falciparum strains. Nucleic Acids Res 34, 1166–1173.

Mackintosh, C.L., Christodoulou, Z., Mwangi, T.W., Kortok, M., Pinches, R., Williams, T.N., Marsh, K., and Newbold, C.I. (2008). Acquisition of naturally occurring antibody responses to recombinant protein domains of Plasmodium falciparum erythrocyte membrane protein 1. Malaria Journal 7, 1–10.

Makobongo, M.O., Keegan, B., Long, C.A., and Miller, L.H. (2006). Immunization of Aotus monkeys with recombinant cysteine-rich interdomain region 1 alpha protects against severe disease during Plasmodium falciparum reinfection. The Journal of infectious diseases 193, 731–740.

Manichaikul, A., Mychaleckyj, J.C., Rich, S.S., Daly, K., Sale, M., and Chen, W.-M. (2010). Robust relationship inference in genome-wide association studies. Bioinformatics 26, 2867–2873.

Mi, H., Muruganujan, A., Ebert, D., Huang, X., and Thomas, P.D. (2018). PANTHER version 14: more genomes, a new PANTHER GO-slim and improvements in enrichment analysis tools. Nucleic acids research 47, D419–D426.

Miura, K. (2016). Progress and prospects for blood-stage malaria vaccines. Expert Rev Vaccines 15, 765–781.

Nasamu, A.S., Falla, A., Pasaje, C.F.A., Wall, B.A., Wagner, J.C., Ganesan, S.M., Goldfless, S.J., and Niles, J.C. (2019). An integrated platform for genome engineering and gene expression perturbation in Plasmodium falciparum. BioRxiv, 816504.

Nkrumah, L.J., Muhle, R.A., Moura, P.A., Ghosh, P., Hatfull, G.F., Jacobs, W.R., Jr., and Fidock, D.A. (2006). Efficient site-specific integration in Plasmodium falciparum chromosomes mediated by mycobacteriophage Bxb1 integrase. Nature methods 3, 615–621.

Oduola, A.M., Milhous, W.K., Weatherly, N.F., Bowdre, J.H., and Desjardins, R.E. (1988). Plasmodium falciparum: induction of resistance to mefloquine in cloned strains by continuous drug exposure in vitro. Experimental parasitology 67, 354–360.

Omelianczyk, R.I., Loh, H.P., Chew, M., Hoo, R., Baumgarten, S., Renia, L., Chen, J., and Preiser, P. (2020). Rapid activation of distinct members of multigene families in Plasmodium sp. Communications Biology Accepted.

Patel, S.N., Serghides, L., Smith, T.G., Febbraio, M., Silverstein, R.L., Kurtz, T.W., Pravenec, M., and Kain, K.C. (2004). CD36 mediates the phagocytosis of Plasmodium falciparum–Infected erythrocytes by rodent macrophages. Journal of Infectious Diseases 189, 204–213.

Peters, J.M., Fowler, E.V., Krause, D.R., Cheng, Q., and Gatton, M.L. (2007). Differential changes in Plasmodium falciparum var transcription during adaptation to culture. The Journal of infectious diseases 195, 748–755.

Petter, M., Haeggström, M., Khattab, A., Fernandez, V., Klinkert, M.-Q., Wahlgren, M.J.M., and parasitology, b. (2007). Variant proteins of the Plasmodium falciparum RIFIN family show distinct subcellular localization and developmental expression patterns. 156, 51–61.

Rénia, L., and Goh, Y.S. (2016). Malaria Parasites: The Great Escape. 7.

Ritchie, M.E., Phipson, B., Wu, D., Hu, Y., Law, C.W., Shi, W., and Smyth, G.K. (2015). limma powers differential expression analyses for RNA-sequencing and microarray studies. Nucleic Acids Res 43, e47.

Rottmann, M., Lavstsen, T., Mugasa, J.P., Kaestli, M., Jensen, A.T.R., Müller, D., Theander, T., and Beck, H.-P. (2006). Differential Expression of var Gene Groups Is Associated with Morbidity Caused by Plasmodium falciparum Infection in Tanzanian Children. Infection and immunity 74, 3904.

Saito, F., Hirayasu, K., Satoh, T., Wang, C.W., Lusingu, J., Arimori, T., Shida, K., Palacpac, N.M.Q., Itagaki, S., Iwanaga, S., et al. (2017). Immune evasion of Plasmodium falciparum by RIFIN via inhibitory receptors. Nature 552, 101–105.

Salanti, A., Staalsoe, T., Lavstsen, T., Jensen, A.T., Sowa, M.K., Arnot, D.E., Hviid, L., and Theander, T.G.J.M.m. (2003). Selective upregulation of a single distinctly structured var gene in chondroitin sulphate A-adhering Plasmodium falciparum involved in pregnancy-associated malaria. 49, 179–191.

Serghides, L., Patel, S.N., Ayi, K., and Kain, K.C. (2006). Placental Chondroitin Sulfate A–Binding Malarial Isolates Evade Innate Phagocytic Clearance. Journal of Infectious Diseases 194, 133–139.

Sidik, S.M., Huet, D., Ganesan, S.M., Huynh, M.H., Wang, T., Nasamu, A.S., Thiru, P., Saeij, J.P.J., Carruthers, V.B., Niles, J.C., et al. (2016). A Genome-wide CRISPR Screen in Toxoplasma Identifies Essential Apicomplexan Genes. Cell 166, 1423-1435.e1412.

Spycher, C., Rug, M., Pachlatko, E., Hanssen, E., Ferguson, D., Cowman, A.F., Tilley, L., and Beck, H.P. (2008). The Maurer’s cleft protein MAHRP1 is essential for trafficking of PfEMP1 to the surface of Plasmodium falciparum-infected erythrocytes. Molecular microbiology 68, 1300–1314.

Theander, T.G., and Lusingu, J.P.A. (2015). Efficacy and safety of RTS, S/AS01 malaria vaccine with or without a booster dose in infants and children in Africa: final results of a phase 3, individually randomised, controlled trial. Lancet 386, 31–45.

Tusher, V.G., Tibshirani, R., and Chu, G. (2001). Significance analysis of microarrays applied to the ionizing radiation response. Proceedings of the National Academy of Sciences 98, 5116–5121.

Udeinya, I.J., Graves, P.M., Carter, R., Aikawa, M., and Miller, L.H. (1983). Plasmodium falciparum: effect of time in continuous culture on binding to human endothelial cells and amelanotic melanoma cells. Experimental parasitology 56, 207–214.

Urban, B.C., Ing, R., and Stevenson, M.M. (2005). Early interactions between blood-stage plasmodium parasites and the immune system. Current topics in microbiology and immunology 297, 25–70.

Voss, T.S., Healer, J., Marty, A.J., Duffy, M.F., Thompson, J.K., Beeson, J.G., Reeder, J.C., Crabb, B.S., and Cowman, A.F. (2006). A var gene promoter controls allelic exclusion of virulence genes in Plasmodium falciparum malaria. Nature 439, 1004–1008.

Walliker, D., Quakyi, I.A., Wellems, T.E., McCutchan, T.F., Szarfman, A., London, W.T., Corcoran, L.M., Burkot, T.R., and Carter, R. (1987). Genetic analysis of the human malaria parasite Plasmodium falciparum. Science (New York, NY) 236, 1661–1666.

Yang, X., Xu, S., and Consortium, H.P.-A.S. (2011). Identification of close relatives in the HUGO Pan-Asian SNP Database. PloS one 6, e29502.

Ye, W., Chew, M., Hou, J., Lai, F., Leopold, S.J., Loo, H.L., Ghose, A., Dutta, A.K., Chen, Q., Ooi, E.E., et al. (2018). Microvesicles from malaria-infected red blood cells activate natural killer cells via MDA5 pathway. PLOS Pathogens 14, e1007298.

Yeda, R., Ingasia, L.A., Cheruiyot, A.C., Okudo, C., Chebon, L.J., Cheruiyot, J., Akala, H.M., and Kamau, E. (2016). The genotypic and phenotypic stability of Plasmodium falciparum field isolates in continuous in vitro culture. Plos one 11, e0143565.

Zenonos, Z.A., Dummler, S.K., Müller-Sienerth, N., Chen, J., Preiser, P.R., Rayner, J.C., and Wright, G.J. (2015). Basigin is a druggable target for host-oriented antimalarial interventions. The Journal of experimental medicine 212, 1145–1151.

